# Genome wide association study reveals plant loci controlling heritability of the rhizosphere microbiome

**DOI:** 10.1101/2020.02.21.960377

**Authors:** Siwen Deng, Daniel Caddell, Jinliang Yang, Lindsay Dahlen, Lorenzo Washington, Devin Coleman-Derr

**Affiliations:** Department of Plant and Microbial Biology, University of California, Berkeley, CA, USA; Plant Gene Expression Center, USDA-ARS, Albany, CA, USA; Department of Agronomy and Horticulture, University of Nebraska-Lincoln, Lincoln, NE, USA; Center for Plant Science Innovation, University of Nebraska-Lincoln, Lincoln, NE, USA; Department of Plant Sciences, University of California, Davis, CA, USA

**Keywords:** Rhizosphere, host genetics, microbiome, GWAS, heritability, amplicon sequencing, sorghum

## Abstract

Host genetics has recently been shown to be a driver of plant microbiome composition. However, identifying the underlying genetic loci controlling microbial selection remains challenging. Genome wide association studies (GWAS) represent a potentially powerful, unbiased method to identify microbes sensitive to host genotype, and to connect them with the genetic loci that influence their colonization. Here, we conducted a population-level microbiome analysis of the rhizospheres of 200 sorghum genotypes. Using 16S rRNA amplicon sequencing, we identify rhizosphere-associated bacteria exhibiting heritable associations with plant genotype, and identify significant overlap between these lineages and heritable taxa recently identified in maize. Furthermore, we demonstrate that GWAS can identify host loci that correlate with the abundance of specific subsets of the rhizosphere microbiome. Finally, we demonstrate that these results can be used to predict rhizosphere microbiome structure for an independent panel of sorghum genotypes based solely on knowledge of host genotypic information.

## Introduction

Recent work has shown that root-associated microbial communities are in part shaped by host genetics^1–4^. A study comparing the root microbiomes of a broad range of cereal crops has demonstrated a strong correlation between host genetic differences and microbiome composition^4^, suggesting that a subset of the plant microbiome may be influenced by host genotype across a range of plant hosts. In maize, these genotype-sensitive, or “heritable”, microbes are phylogenetically clustered within specific taxonomic groups^5^; however, it is unclear whether the increased genotype sensitivity in these lineages is unique to the maize microbiome or is common to other plant hosts as well.

Despite consistent evidence of the interaction between host genetics and plant microbiome composition, identifying specific genetic elements driving host-genotype dependent microbiome acquisition and assembly in plants remains a challenge. Recent efforts guided by *a priori* hypotheses of gene involvement have begun to dissect the impact of individual genes on microbiome composition^6,7^. However, these studies are limited to a small fraction of plant genes predicted to function in microbiome-related processes. Additionally, many plant traits expected to impact microbiome composition and activity, such as root exudation^8^ and root system architecture^9^, are inherently complex and potentially governed by a very large number of genes. For these reasons, there is a need for alternative, large-scale and unbiased methods for identifying the genes that regulate host-mediated selection of the microbiome.

Genome-wide association studies (GWAS) represent a powerful approach to map loci that are associated with complex traits in a genetically diverse population. Though pioneered for use in human genetics, to date the majority of GWAS have been conducted in plants^10^, and it has become an increasingly popular tool for studying the genetic basis of natural variation and traits of agricultural importance. When inbred lines are available, GWAS can be particularly useful; once genotyped, these lines can be phenotyped multiple times, making it possible to study many different traits in many different environments^11^. While GWAS is typically used in the context of a single quantitative phenotypic trait, analyses of multivariate molecular traits, such as transcriptomic or metabolomic data, have also been conducted^12,13^. More recently, several attempts have been made to use host-associated microbiome census data as an input to GWAS, which in theory will allow for the identification of host genetic loci controlling microbiome composition^14,15^.

In plants, a recent study in *Arabidopsis thaliana* used phyllosphere microbial community data as the phenotypic trait in a GWAS to demonstrate that plant loci responsible for defense and cell wall integrity affect microbial community variation^16^. Several other recent phyllosphere studies performed GWAS to identify genetic factors controlling microbiome associations with mixed degrees of success^16–18^. However, to our knowledge, use of GWAS in conjunction with the root associated microbiome has yet to be explored. In the context of the root microbiome, selection of sample type (rhizosphere or endosphere) and host system may be critical factors that determine the success of such effort. Previous work comparing the root microbiomes of diverse cereal crops have offered conflicting evidence as to whether host genotypic distance correlates most strongly with microbial communities distance within root endospheres or rhizospheres^3,4^. These data suggest that the sample type exhibiting the strongest correlation between genotype and microbiome composition may differ for each host, and that an initial evaluation of the degree of correlation between genotype and microbiome phenotype across sample types may be informative.

In the context of the root microbiome, we propose *Sorghum bicolor* (L.) as an ideal plant system for GWAS-based dissection of host-genetic control of microbiome composition. Sorghum is a heavy producer of root exudates^19^, and the sorghum microbiome has been shown to house an unusually large number of host-specific microbes^4^. Additionally, there is a wide range of natural adaptation in traditional sorghum varieties from across Africa and Asia, and a collection of breeding lines generated from U.S. sorghum breeding programs, both of which provide a rich source of phenotypic and genotypic variation^20^. Several genome sequences of sorghum varieties have been completed, and variation in nucleotide diversity, linkage disequilibrium, and recombination rates across the genome have been quantified^21^, providing an understanding of the genomic patterns of diversification in sorghum. Finally, sorghum is an important cereal crop grown throughout the world as a food, feedstock, and biofuel, enabling direct integration of resulting discoveries into an agriculturally-relevant system.

In this study, we dissect the host-genetic control of bacterial microbiome composition in the sorghum rhizosphere. Using 16S rRNA sequencing, we profiled the microbiome of a panel of 200 diverse genotypes of field grown sorghum. We aim to demonstrate that a large fraction of the plant microbiome responds to host genotype, and that this subset shares considerable overlap with lineages shown to be susceptible to host genetic control in another plant host. Additionally, we tested whether GWAS can be used to identify specific genetic loci within the host genome that are correlated with the abundance of specific heritable lineages, and whether differences in microbiome composition can be predicted solely from genotypic information. Collectively, this work demonstrates the utility of GWAS for analysis of host-mediated control of rhizosphere microbiome phenotypes.

## Results

### Diverse sorghum germplasm show rhizosphere is ideal for microbiome-based GWAS

In this study, the relationship between host genotype and microbiome composition was explored through a field experiment involving 200 genotypes selected from the Sorghum Association Panel (SAP) germplasm collection^20^ (Supplemental Table 1). As prior studies suggest that the strength of the correlation between host genotype and microbiome composition may vary by sample type in a host-dependent manner ^3,4^, we first sought to determine whether leaf, root, or rhizosphere samples were most suitable for downstream GWAS in sorghum. Using a subset of 24 genotypes from our collection of 200 (Figure 1a, Supplemental Table 1), the microbiome composition of leaf, root, and rhizosphere sample types was analyzed using paired-end sequencing of the V3–V4 region of the ribosomal 16S rRNA on the Illumina MiSeq platform (Illumina Inc., San Diego, CA, USA). The resulting dataset demonstrated comparatively high levels of microbial diversity within both root and rhizosphere samples (Figure 1b) and strong clustering of above and below ground sample types (Figure 1c). Three independent Mantel’s tests (9,999 permutations) were used to evaluate the degree of correlation between host genotypic distance and microbiome composition for leaf, root, and rhizosphere sample types (Figure 1d); of the three compartments, only rhizosphere exhibited a significant Mantel’s correlation (R^2^=0.13, Df=1, p=0.02). Based on these results, subsequent investigation of the microbiomes of the full panel of 200 lines, including heritability and GWAS analyses, was performed using rhizosphere samples.

**Figure 1.**
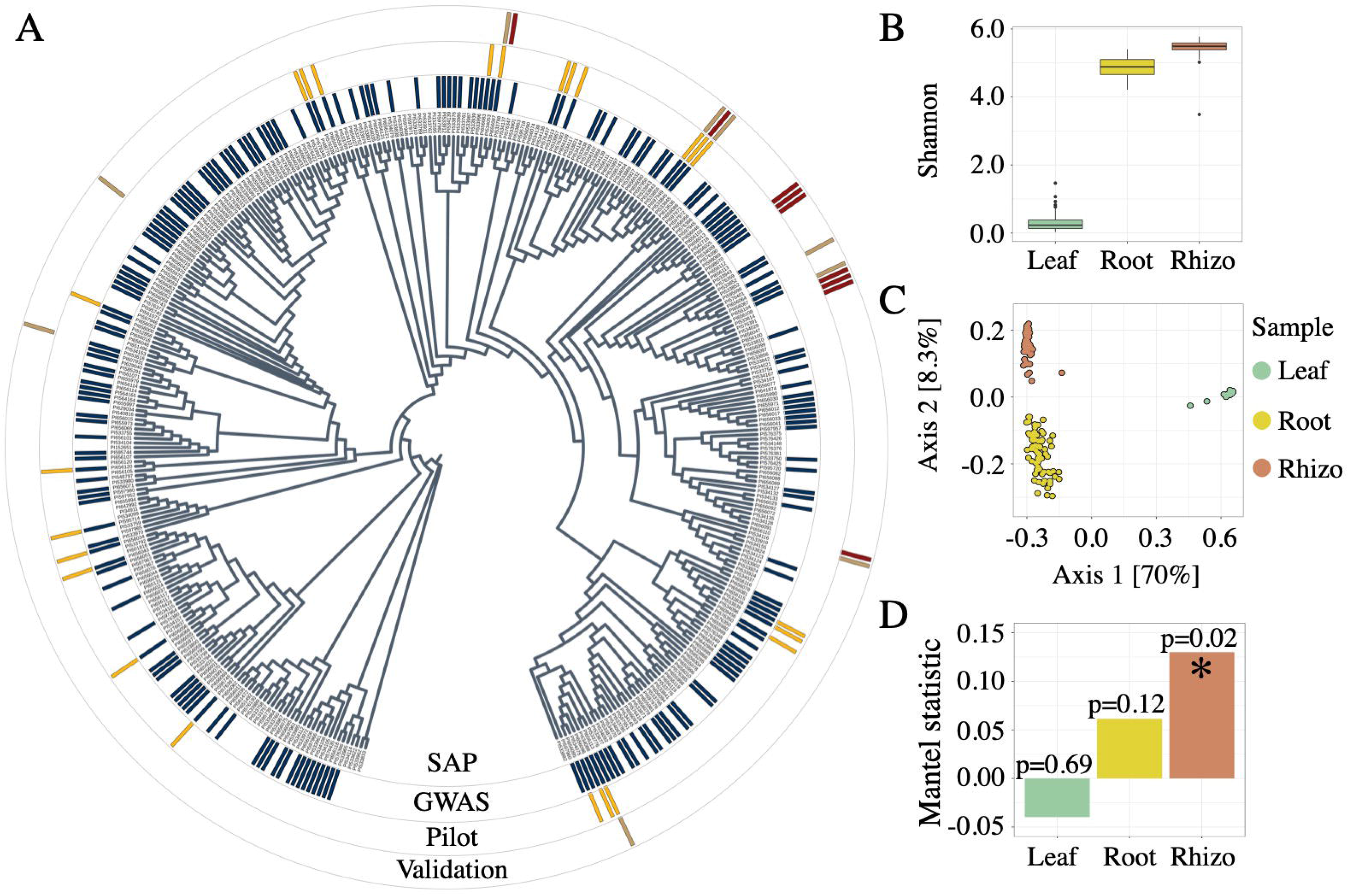
Sample type and population selection. **A** Phylogenetic tree representing the 378 member sorghum association panel (SAP, inner ring), the subset of 200 lines selected for GWAS (2nd ring from the center, in blue), the 24 lines used for sample type selection (Pilot, 3rd ring from the center, in yellow), and the 18 genotypes used for GWAS validation containing either the Chromosome 4 minor allele (red) or major allele (brown) identified by GWAS (outer ring). **B** Shannon’s Diversity values from 16S rRNA amplicon datasets for the leaf (green), root (yellow), and rhizosphere (red) sample types across all 24 genotypes used in the pilot experiment. **C** Principal coordinate analysis generated using Bray-Curtis distance for the 24 genotypes across leaf (green), root (yellow), and rhizosphere (red). **D** Mantel’s R statistic plotted for each sample type indicating the degree of correlation between host genotypic distance and microbiome distance.

To investigate host genotype dependent variation in the sorghum rhizosphere microbiome, the rhizospheres of 600 field grown plants (including three replicates of each of 200 genotypes) were profiled using V3-V4 16S rRNA amplicon sequencing. After removing rare OTUs with less than 3 reads in at least 20% of the samples and normalizing to an even read depth of 18,000 reads per sample, the data set included 1,189 high-abundance OTUs representing 29 bacterial phyla. Compositional analysis of the resulting microbiome dataset exhibited profiles consistent with recent microbiome studies involving the sorghum rhizosphere^4,22,23^ from a variety of field sites, with Proteobacteria, Actinobacteria and Acidobacteria comprising the top three dominant phyla (Supplemental Figure 1).

### Sorghum and maize rhizospheres exhibit strong overlap in heritable taxa

A recent study of two separate maize microbiome datasets suggests that specific bacterial lineages are more sensitive to the effect of host genotype than others^5^. To determine if a bacterial lineage’s responsiveness to host genetics is a trait conserved across different plant hosts that diverged more than 11 million years ago^24^, the broad sense heritability (H^2^) of individual OTUs in our sorghum dataset was evaluated. H^2^, which quantifies the proportion of variance that is explained by genetic rather than environmental effects, ranged from 0 to 66% for individual OTUs (Supplemental Table 2). By comparison, H^2^ for individual OTUs in the first of two experiments across 27 inbred maize lines had a maximum of 23% (performed in 2010), while the second exhibited a maximum of 54% (performed in 2015)^5^.

To explore whether microbes with high heritability in the sorghum dataset are phylogenetically clustered, we partitioned the 1,189 OTUs into heritable (n=347) and non-heritable fractions (n=842) using an H^2^ cutoff score of 0.15 (Figure 2a, Supplemental Table 3). Several bacterial orders, including Verrucomicrobiales, Flavobacteriales, Planctomycetales, and Burkholderiales, were observed to have significantly greater numbers of OTUs that are heritable, as compared to the non-heritable OTU fraction (Fisher’s exact test, p<0.05, Figure 2a, Supplemental Table 3). Notably, all 6 Flavobacteriales OTUs were present in the heritable fraction (Figure 2b); by contrast, 40 other bacterial orders were only observed within the non-heritable fraction. Another bacterial order, Bacillalles, contained a smaller number of OTUs in the heritable than non-heritable fraction, but the percentage of read counts attributable to its heritable OTUs was approximately eight-fold greater than those in the non-heritable fraction, suggesting that its heritable members are abundant organisms within the rhizosphere (Figure 2b). Collectively, these data demonstrate that a specific subset of bacterial lineages are enriched for members susceptible to host genotypic selection.

**Figure 2.**
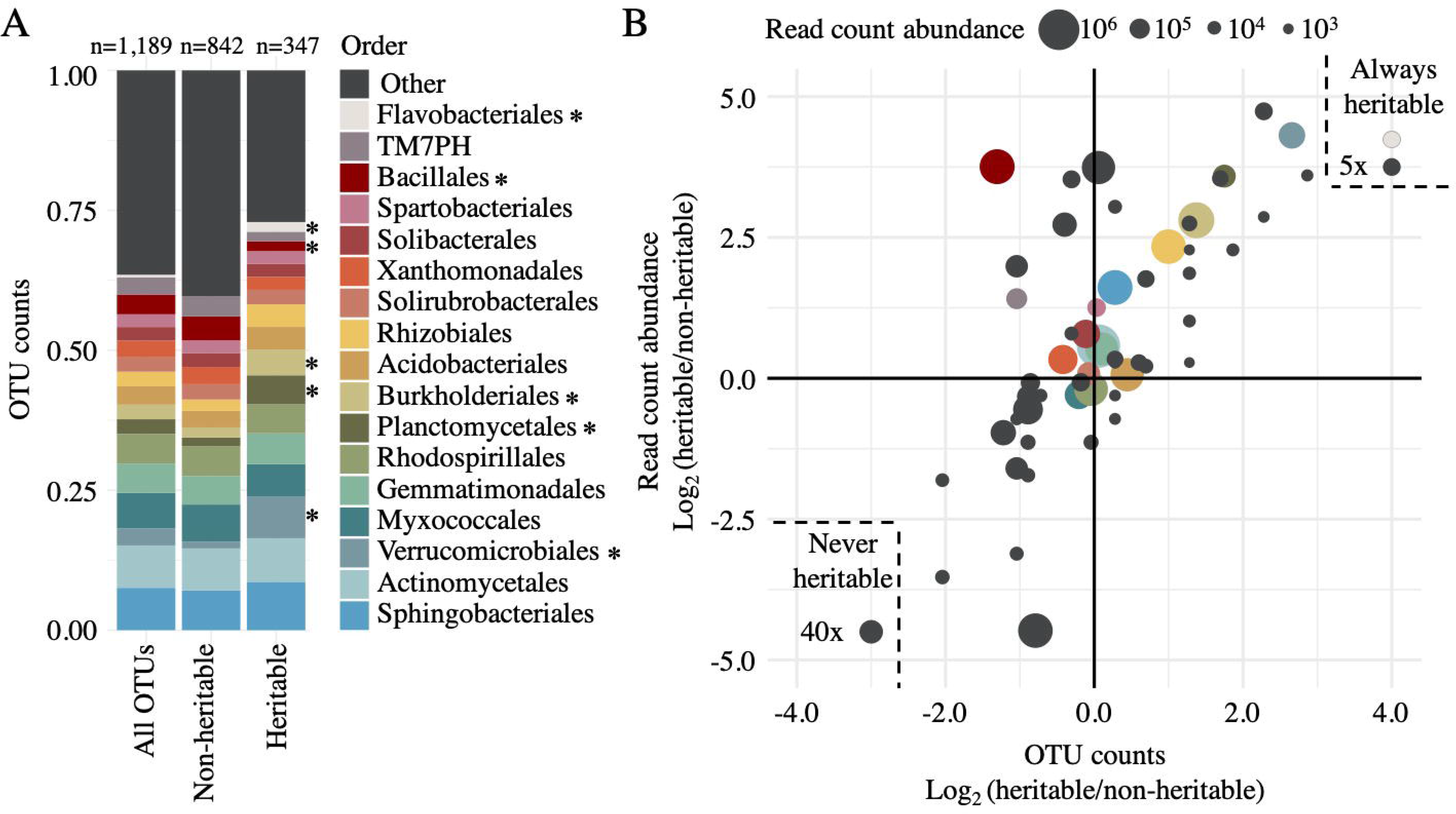
Taxonomic classification of heritable rhizosphere microbes. **A** The relative percentage of total OTUs belonging to each of the top 17 bacterial orders for all OTUs (left bar), non-heritable OTUs (middle bar), or heritable OTUs (right bar). Orders with significantly different numbers of OTUs in the heritable (H^2^>0.15) as compared to the non-heritable fraction (H^2^<0.15), as determined by Fisher’s exact test (p<0.05), are indicated with asterisks. **B** Order-level scatterplot of the log2 ratio between heritable and non-heritable OTU counts (x-axis) and read count abundance (y-axis). Circle sizes represent the total abundance represented by each bacterial order. Points within the dashed lines indicate merged bacterial orders that were present only in the heritable (upper right) or non-heritable (lower left) fractions.

We hypothesized that despite the considerable evolutionary distance between maize and sorghum, the bacterial lineages containing OTUs most responsive to host genotypic effects in maize would likely also contain OTUs exhibiting such susceptibility within sorghum. To test this, we compared the top 100 most heritable OTUs from both maize datasets (referred to as NAM 2010 and NAM 2015) and the sorghum dataset described above, resulting in a combined dataset of 300 OTUs spanning 65 bacterial orders. After removing bacterial orders not observed in the sorghum dataset (n=18), we noted that more than half were observed in at least two of the datasets, and approximately one third (n=15) contained heritable OTUs in all three datasets (Figure 3a). To determine if this overlap was significantly greater than is expected by chance, we performed permutational resampling of 10,000 sets of randomly chosen sorghum OTUs for comparison. Notably, we found that the overlap between the heritable sorghum fraction with both the individual maize heritable fractions and the combined heritable maize OTUs to be significant, compared with the resampled sorghum OTUs (NAM 2010 n=17, p=0.0099, NAM 2015 n=19, p=0.0016, combined n=15, p=0.0344)(Figure 3a). Collectively, these results demonstrate that there is a conservation between the bacterial orders most sensitive to genotype across both maize and sorghum.

**Figure 3.**
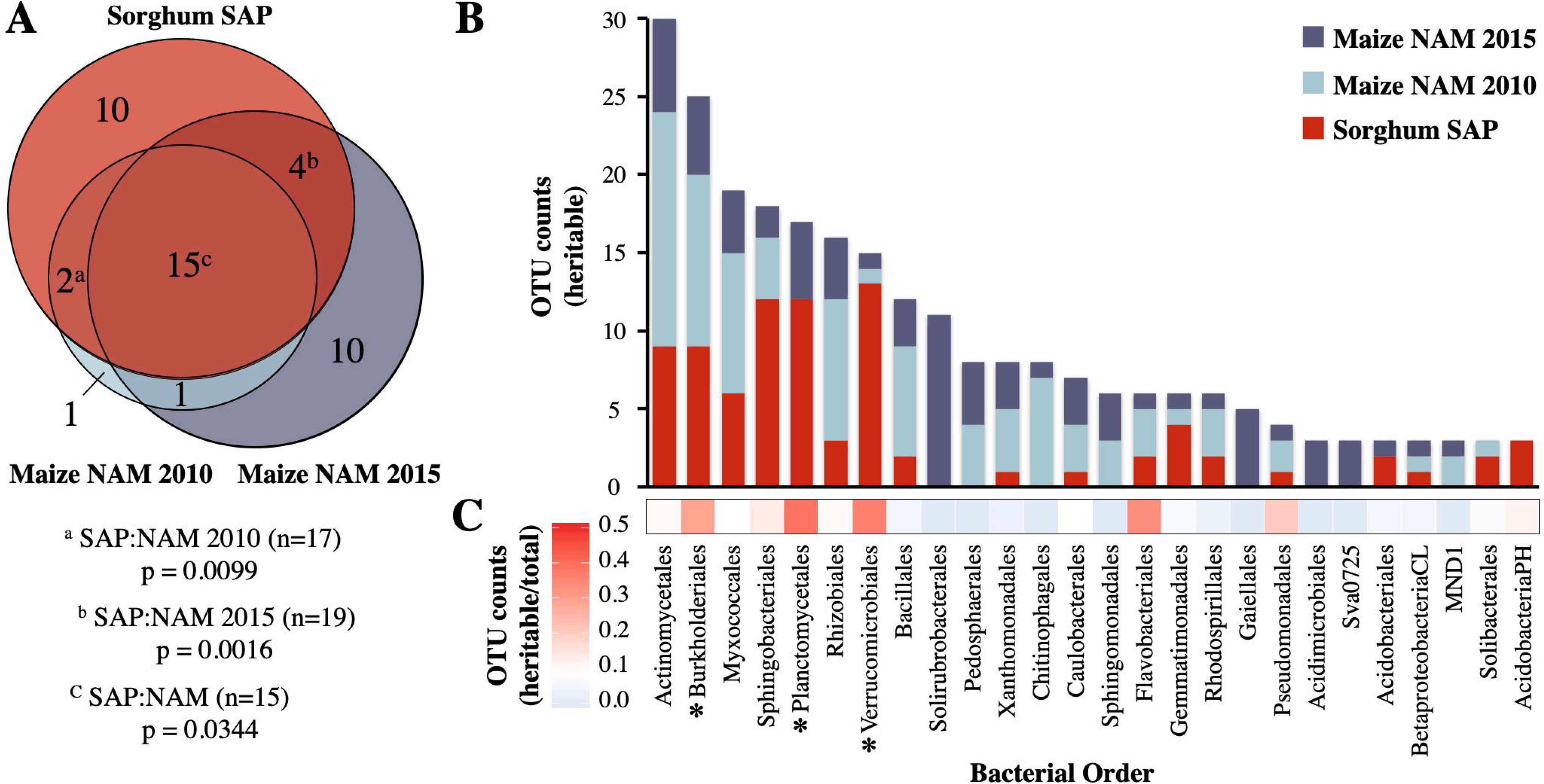
Heritability of rhizosphere microbes across maize and sorghum. **A** Proportional Venn diagram of bacterial orders containing heritable OTUs identified in this study (Sorghum SAP), compared with those found in a large-scale field study of maize nested association mapping (NAM) parental lines grown over two separate years, published in Walters et al., 2018^5^. The top 100 heritable OTUs (based on H^2^) from each dataset were classified at the taxonomic rank of order to generate the Venn diagram. NAM heritable orders only present in the SAP non-heritable fraction are represented by the blue sections. Superscript letters indicate the frequency that a random subsampling of 100 sorghum OTUs (10,000 permutations) produced greater overlap with maize OTUs from either single year (a/b) or both (c). **B** Stacked barplot displaying cumulative counts (y-axis) of OTUs identified as heritable in any of the three datasets for all bacterial orders (x-axis) which have a total of at least three heritable OTUs. **C** The fraction of heritable sorghum OTUs relative to all sorghum OTUs within each order are displayed as a heatmap. Asterisks indicate orders enriched in heritable OTUs (Fisher’s exact test, p<0.001).

In an effort to identify the bacterial lineages with the greatest propensity for high heritability, we calculated the number of heritable OTUs in each of the shared heritable bacterial orders identified above. We noted that among bacterial orders containing the greatest number of heritable OTUs across all three datasets were several that represent large lineages frequently observed within the root microbiome; (e.g. Actinomycetales) (Figure 3b). We hypothesized that this result is likely driven in part by the overall frequency of these lineages within the rhizosphere microbiome, with more common lineages resulting in a greater fraction of heritable microbes due to their ubiquity. To help account for this, we normalized the frequency of heritable sorghum OTUs (n=100) by total sorghum OTU counts (n=1,189) belonging to each order (Figure 3c, Supplemental Table 4). These results demonstrate that while the prevalence of Actinomycetales and Myxococcales among heritable microbes is consistent with their general prevalence in the overall dataset, Burkholderiales and two other lineages, including the Verrucomicrobia and Planctomycetes, exhibited a significant enrichment (Fisher’s exact test, p<0.001) in the heritable fraction not expected to be influenced by abundance alone.

### Genome-wide association reveals genetic loci correlated with rhizosphere microbial abundance

Recent work in the leaf microbiome has demonstrated the potential utility of GWAS for uncovering host loci correlated with microbiome composition^18^. Here, we sought to use GWAS with rhizosphere microbiome datasets using both global properties of the OTU dataset and the abundances of individual OTUs. For overall community composition, a subset of principal components (PCs) were selected from an analysis of the abundance patterns of the 1,189 OTUs. To prioritize individual PCs for inclusion in our GWAS analysis, we determined the heritability scores of each of the top ten PCs, which explained 75% of the total variance in our dataset (Supplemental Figure 2a). PCs with H^2^ equal to or greater than 0.25 (PC1, PC3, PC5, PC9, and PC10, Supplemental Figure 2a) were subjected to GWAS (Supplemental Figure 2b). The GWAS analysis performed for PC1, which explained 21% percent of total variance and had the second highest heritability (H^2^=0.35), revealed a significant correlation between community composition and a locus of approximately 1.15 Mb on chromosome 4 with a moderately stringent threshold of −log_10_ (p=10^−4^) (Figure 4a, Supplemental Figure 2b). Additionally, GWAS analyses that used PC5 and PC10 as inputs, both revealed an identifiable peak on chromosome 6, though it was slightly below the threshold of significance (Supplemental Figure 2b).

**Figure 4.**
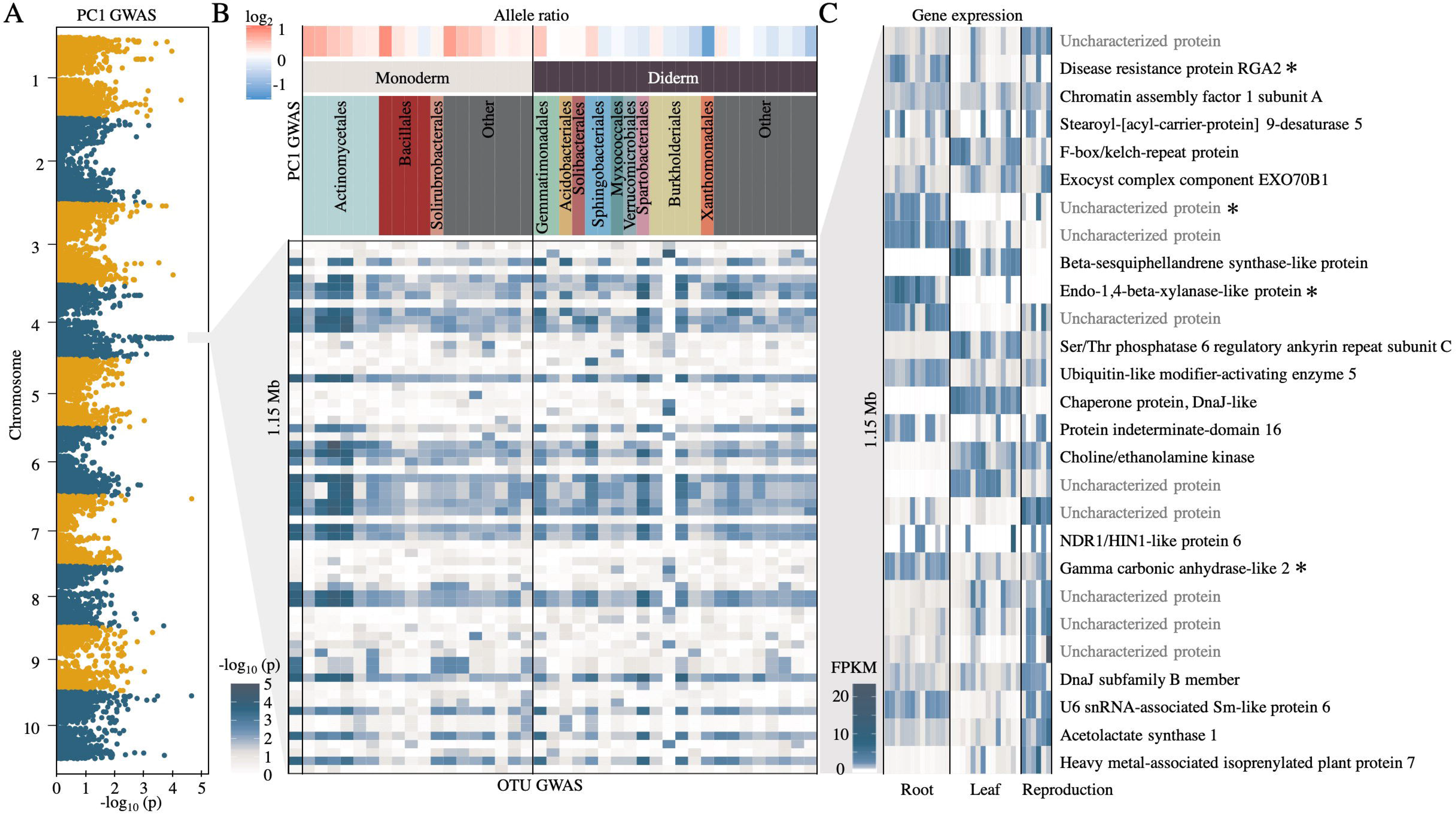
A sorghum genetic locus is correlated with rhizosphere microbial abundance. **A** Manhattan plot of PC1 community analysis GWAS. **B** Individual OTU GWAS of all OTUs with at least 5 SNPs above a threshold of −log_10_ (p=10^−2.5^) in the 1.15 Mb window identified on the same chromosome 4 locus identified by PC1 GWAS (lower heatmap). Ratio of OTUs that associate with the sorghum major (red) or minor (blue) allele groups within this locus (upper heat map). OTUs were grouped based on the predicted presence of one or two membranes (monoderm or diderm) within each bacterial order and colored as in figure 2. **C** Tissue-specific gene expression data for sorghum genes within the chromosome 4 locus. Darker blue indicates higher expression (normalized FPKM). Asterisks indicate genes whose expression are predicted to be root-specific.

As principal components are derived from linear combinations of the abundance of individual OTUs within the dataset, it is unclear whether the correlations observed on chromosomes 4 and 6 are driven by one common or two different sets of microbial lineages. To address this, we performed separate GWAS analyses using the abundances of each single OTU in our dataset as input (Figure 4b, Supplemental Figure 2c). From these analyses, we identified two distinct sets of 39 and 10 OTUs with significant correlations with the loci on chromosomes 4 and 6, respectively, and only a single OTU belonging the the order Burkholderiales that was shared between the two loci (Supplemental Figure 2c), demonstrating that different sorghum loci influence the abundance patterns of different groups of microbes.

To explore the relationship between the identified peak on chromosome 4 (Figure 4a) and the bacterial taxa with significant GWAS correlations at this locus (Figure 4b), we first sought to understand how relative abundance for these 40 OTUs varied across the sorghum panel. An analysis of the SNP data at this locus revealed two allele groups, the major allele containing 343 sorghum genotypes and the minor allele containing 14 genotypes. Next, we observed that the majority of OTUs that were more prevalent in sorghum genotypes containing the major allele belonged to monoderm lineages, while the majority of OTUs more prevalent in the minor allele group belonged to diderm lineages (Figure 4b), suggesting that host genetic mechanisms at this locus are interacting with basal bacterial traits.

To explore which genetic mechanisms might be driving the correlations observed on Chromosome 4, we examined tissue specific expression patterns from publicly available RNA-Seq datasets obtained from phytozome v12.1^25^ for all 27 genes in the 1.15 Mb interval (Figure 4c, Supplemental Table 5). Of these candidates, several were observed to exhibit strong root specific expression patterns, including three annotated candidates: gamma carbonic anhydrase-like 2, a putative Beta-1,4 endoxylanase, and disease resistance protein RGA2 (Figure 4c).

### Sorghum genotypic data can predict microbiome composition

To validate that allelic variation at the candidate locus on chromosome 4 contributes to differences in rhizosphere composition, we conducted a follow up experiment with eighteen additional sorghum lines, including genotypes not present in the original study. To help disentangle phylogenetic-relatedness from locus-specific effects, we selected sorghum genotypes that spanned the diversity panel; additionally, for each minor allele genotype (n=9), we included a phylogenetically related major allele line (n=9) (Figure 1a). Following two weeks of growth in a mixture of calcined clay and field soil in the growth chamber, we collected the rhizosphere microbiomes of each genotype and microbiome composition was analyzed using 16S rRNA amplicon sequencing as in the main study. A canonical analysis of principal coordinates (CAP) ordination constrained on genotypic group separated the rhizospheres of genotypes belonging to major and minor allele groups into distinct clusters (Figure 5a, PERMAnova F=2.66, Df=1, p=0.0061), with genotype explaining approximately 7.5% (CAP1) of variance in the dataset.

**Figure 5.**
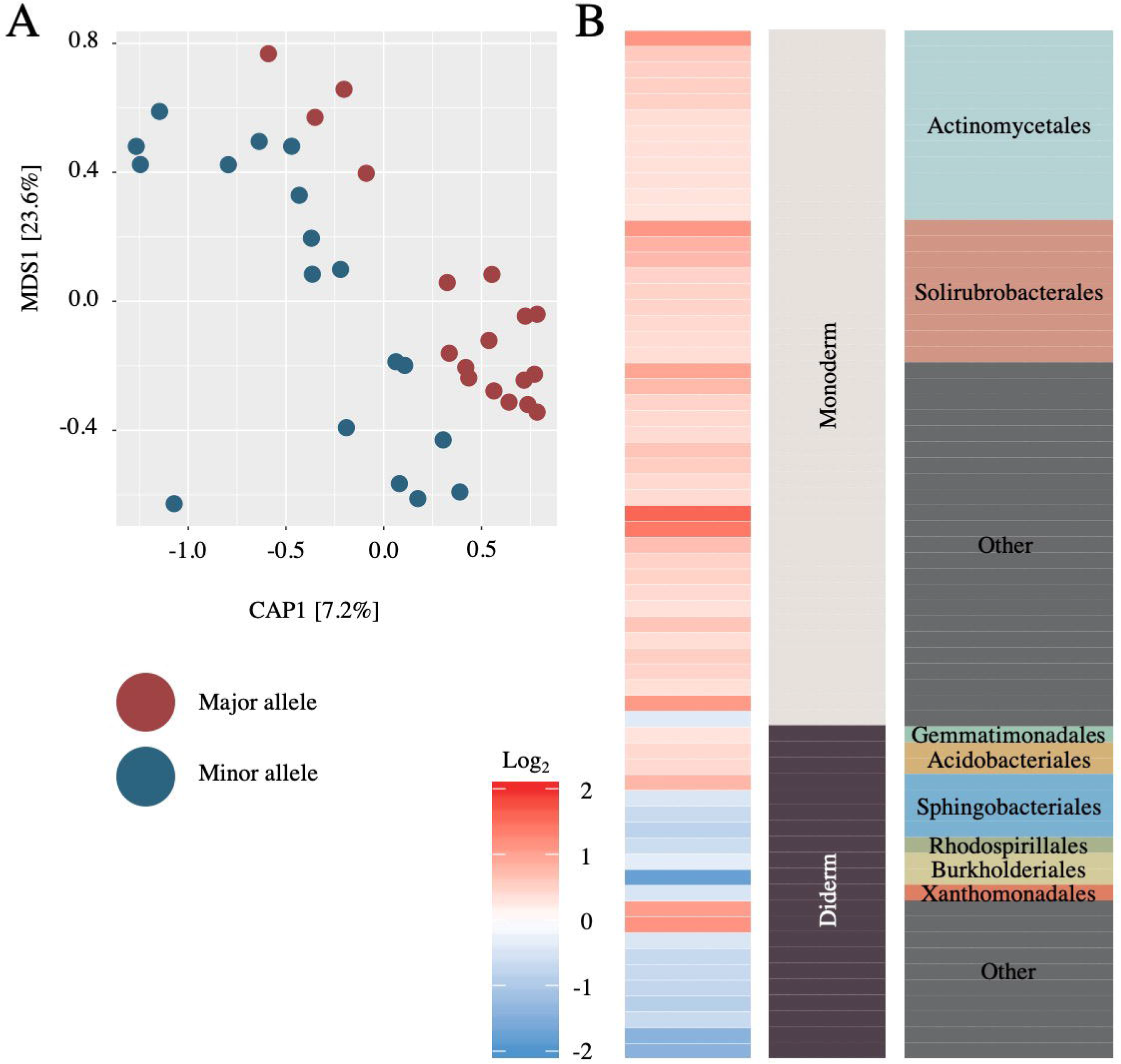
Sorghum genetic information can be used to predict rhizosphere microbiome composition. **A** Canonical Analysis of Principal Coordinates of the rhizosphere microbiome for nine major allele genotypes (red) and nine minor allele genotypes (blue). **B** Ratio of indicator OTUs that associate with the sorghum major (red) or minor (blue) allele groups. OTUs were grouped based on the predicted presence of one or two membranes (monoderm or diderm), within each bacterial order, and colored as in figures 2 and 4.

To identify which taxa drive the clustering observed in our CAPs analysis, and to compare this to taxa responsive to the chromosome 4 allele group in our main experiment, we performed an indicator species analysis on the validation dataset. A comparison of the significant indicator OTUs (p<0.05) from each allele group in the validation dataset (n=65) demonstrated similar trends in abundance of indicator OTUs as observed in the main experiment (Figure 4b), with OTUs belonging to monoderm and diderm lineages enriched in the major and minor allele-containing lines, respectively. Interestingly, while most diderm lineages were more prevalent in the minor allele-containing lines, several diderm lineages including Gemmatimonadales, Acidobacteriales, and Sphingobacteriales contained OTUs that were more abundant within major allele lines. Notably, this pattern was observed in both the main experiment (Figure 4b) and validation experiment (Figure 5b). Collectively, this experiment supports the findings of our main experiment, in which allelic variation at a locus located on chromosome 4 was shown to correlate with the abundance of specific bacterial lineages.

## Discussion

### Host selection of plant rhizosphere microbiomes

Previous GWAS of plant-associated microbiome traits have often been conducted with leaf samples, and have not always been successful in identifying loci that correlate with microbiome phenotypes^16–18^. In this study, we compared the overall correlation between host genotype and bacterial microbiome distances across leaf, root, and rhizosphere of *Sorghum bicolor*, and demonstrate that of the three, the rhizosphere represents the most promising compartment for conducting experiments to untangle the heritability of the sorghum microbiome. Notably, the degree of correlation between sorghum phylogenetic distance and microbiome distance was highest in the rhizosphere and lowest in the leaves. This greater correlation observed in the rhizosphere could be in part due to the phyllosphere’s relative compositional simplicity. Even *Arabidopsis* rosette leaves, which are in close proximity to soil, harbor a distinct and relatively simple bacterial community compared to the root^26^. By contrast, the rhizosphere represents a highly diverse and populated subset of the soil microbiome, and potentially offers a greater pool of microbes upon which the host may exert influence^27^. Alternatively, the rhizosphere’s greater correlation with microbiome composition could be caused by the plant’s relatively weaker ability to select epiphytes in its aboveground microbiome; while the arrival of phyllosphere colonists is largely thought to be driven by wind and rainfall dispersal^28^, root exudation is known to control chemotaxis and other colonization activities of select members of the surrounding soil environment. This provides an additional mechanism for host selection of its microbial inhabitants prior to direct interaction with the plant surface^8,29,30^. It is worth noting that sorghum is known to be an atypically strong producer of root exudates^19^, and consequently it is possible that other plant hosts may demonstrate the greatest selective influence within tissues other than the rhizosphere. Future efforts to investigate host control of the microbiome through GWAS or related techniques would benefit from careful selection of sample type following pilot studies designed to explore heritability across different host tissues.

### Heritable rhizosphere microbes are phylogenetically clustered and similar across hosts

Within the rhizosphere, we demonstrate that microbiome constituents vary in broad sense heritability, and heritable taxa show a strong overlap with heritable lineages identified in maize, spanning fifteen different bacterial orders^5^. In particular, three of these orders, Verrucomicrobiales, Burkholderiales, and Planctomycetales were significantly enriched in the heritable fraction of our dataset. As members of Burkholderiales can form symbioses with both plant and animal hosts^31,32^, and some colonize specific members of a host genus or species^33^, it is feasible that such strong relationships necesitated additional genetic discrimination between hosts. Within *Burkholderia* spp., this could be facilitated by their relatively large pan-genome, with diversity driven by large multi-replicon genomes and abundant genomic islands ^34^.

These observations suggest that evaluating bacterial heritability may identify new lineages for which close or symbiotic but previously undetected associations with plant hosts exist. For example, we observed several lineages with high heritability that are common in soil, yet prior evidence of plant-microbe interactions in the literature is lacking, including Verrucomicrobiales and Planctomycetales. Interestingly, heritability in these lineages might be facilitated by the presence of a recently discovered shared bacterial microcompartment gene cluster present in both Planctomycetes and Verrucomicrobia, which confers the ability to degrade certain plant polysaccharides^35^. Indeed, microbiome composition is known to be driven in part by variations in polysaccharide containing sources including plant cell wall components and root exudates^36^. Additional experimentation with bacterial mutants lacking this genetic cluster could be useful for revealing its role in shaping plant microbe interactions.

### Sorghum loci are responsible for controlling the rhizobiome

Our GWAS correlated host genetic loci and the abundance of specific bacteria within the host microbiome, as well as overall rhizosphere community structure. To our knowledge, this is the first example of such work in a crop rhizosphere. We identified two loci with strong associations with the microbiome structure. The most significant maps to a locus on chromosome 4 containing several candidate genes with root specific expression.

One candidate gene located near the center of this locus encodes a beta 1,4 endo xylanase. Xylanases are responsible for the degradation of xylan into xylose, and are one of the primary catabolizers of hemicellulose, a major component of the plant cell wall^37^. As a result, beta 1,4 endo xylanases may play a role in shaping the degree of plasticity in the barrier between the root and surrounding rhizosphere environments, in turn influencing the release of cell wall or apoplast derived metabolites into the rhizosphere environment^38^. Alternatively, altered xylanase activity could lead to shifts in carbohydrate profiles within the cell wall, leading to heightened plant immune responses^39,40^; the catabolic byproducts of microbially-produced xylanase used in pathogen invasion are in part responsible for triggering innate immune responses in plants, and various components of the plant immune signalling network have been shown to influence microbiome structure^6,7^.

Another candidate gene within the chromosome 4 locus, that also displays root-specific expression, is predicted to encode gamma carbonic anhydrase-like 2. In plants, carbonic anhydrases (CA) participate in aerobic respiration, and facilitate the reversible hydration of CO2 to bicarbonate^41,42^. Previous studies have implicated CA activity in plant-microbe interactions^43^; an important role for CA was first observed in root nodules of legumes inoculated with Rhizobium^44,45^. CAs have since been implicated in disease resistance as well, having both antioxidant activity and salicylic acid binding capability^46–48^. Collectively, these studies suggest that a loss or alteration of function of CA could impact the composition of the rhizosphere microbiome. Future validation experiments using genetic mutants within this and other candidate genes can be used to help elucidate the underlying genetic element(s) responsible for modulation of the rhizosphere microbiome.

## Conclusion

Although the underlying host genetic causes of shifts in the microbiome are not well understood, candidate driven approaches have implicated disease resistance^6,7^, nutrient status^7,49,50^, sugar signaling^51^, and plant age^52,53^ as major factors. Non-candidate approaches to link host genetics and microbiome composition, such as GWAS, have the potential to discover novel mechanisms that can be added to this list. Here we show that GWAS can predict microbiome structure based on host genetic information, building on previous studies that have observed inter- and intra-species variation in microbiomes^1,4,5,16,36,54–56^. Collectively, our study adds to a growing list of evidence that genetic variation within plant host genomes modulates their associated microbiome. We anticipate that GWAS of plant microbiome association will promote a comprehensive understanding of the host molecular mechanisms underlying the assembly of microbiomes and facilitate breeding efforts to promote beneficial microbiomes and improve plant yield.

## Methods

### Germplasm selection

In order to ensure that microbiome profiling was performed on a representative subset of the broad genetic diversity present in the 378 member Sorghum Association Panel (SAP)^20^, subsets of 200 genotypes were randomly sampled from the SAP 10,000 times and an aggregate nucleotide diversity score was calculated for each using the R package “PopGenome”^57^. From these data, the subset of 200 lines with the maximum diversity value was selected (Figure 1a, Supplemental Table 1). For the pilot experiment used to determine the appropriate sample type for GWAS, a subset of 24 lines was selected that included genotypes from a wide range of phylogenetic distances (Figure 1a, Supplemental Table 1). The phylogenetic tree of sorghum accessions was generated using the online tool: Interactive Tree Of Life (iTOL) v5^58^.

### Field experimental design and root microbiome sample collection

The experimental field used in this study is an agricultural field site located in Albany, California (37.8864°N, 122.2982°W), characterized by a silty loam soil with pH 5.2^4^. Germplasm for the US SAP panel used in this study^20^ were obtained from GRIN (www.ars-grin.gov). To ensure a uniform starting soil microbiome for all sorghum seedlings and to control their planting density, seeds were first sown into a thoroughly homogenized field soil mix in a growth chamber with controlled environmental factors (25 °C, 16hr photoperiods) followed by transplantation to the field site. To prepare the soil for seed germination, 0.54 cubic meters of soil was collected at a depth of 0 to 20 cm from the field site subsequently used for planting, and homogenized by separately mixing 4 equally sized batches with irrigation water in a sterilized cement mixer followed by manual homogenization on a sterilized tarp surface. Soil was then transferred to sterilized 72-cell plant trays. To prepare seeds for planting, seeds were surface-sterilized through soaking 10 min in 10% bleach + 0.1% Tween-20, followed by 4 washes in sterile water. Following planting, sorghum seedlings were watered with approximately 5 ml of water using a mist nozzle every 24 hrs for the first three days, and bottom watered every three days until the 12th day, then transplanted to the field.

The field consisted of three replicate blocks, with each block containing 200 plots for each of 200 selected genotypes. Six healthy sorghum seedlings of each genotype were transplanted to their respective plots, separated by 15.2cm, and thinning to three seedlings per plot was performed at two weeks post transplanting. Plots were organized in an alternating pattern with respect to the irrigation line to maximize the distance between each plant (Supplemental Figure 3). Plants were watered for one hour, three times per week, using drip irrigation with 1.89 L/hour rate flow emitters. Manual weeding was performed three times per week throughout the growing season. To ensure that the genotypes were at a similar stage of development and that the host-associated microbiome had sufficient time to develop, collection of plant-associated samples was performed nine weeks post germination. Only the middle plant within each plot was harvested to help mitigate potential confounding plant-plant interaction effects resulting from contact with roots from neighboring plants of other genotypes. Rhizosphere, leaf, and root samples were collected as described previously^59^.

### DNA extraction, PCR amplification, and Illumina sequencing

DNA extractions, PCR amplification of the V3-V4 region of the 16S rRNA gene, and amplicon pooling were performed as described previously^59^. In brief, DNA extractions for all samples were performed using extraction kits (MoBio PowerSoil DNA Isolation Kit, MoBio Inc., Carlsbad, CA) following the manufacturer’s protocol. Amplification of the V3-V4 region of the 16S rRNA gene was performed using dual-indexed 16s rRNA Illumina iTags primers 341F (5’-CCTACGGGNBGCASCAG-3’) and 785R (5’-GACTACNVGGGTATCTAATCC-3’). An aliquot of the pooled amplicons was diluted to 10 nM in 30μL total volume before submitting to the QB3 Vincent J. Coates Genomics Sequencing Laboratory facility at the University of California, Berkeley for sequencing using Illumina Miseq 300bp pair-end with v3 chemistry. Sequences were returned demultiplexed, with adaptors removed.

### Amplicon sequence processing, OTU classification, and taxonomic assignment

Sequencing data were analyzed using the iTagger pipeline to obtain OTUs^60^. In brief, after filtering 81,416,218 16S rRNA raw reads for known contaminants (Illumina adapter sequence and PhiX), primer sequences were trimmed from the 5’ ends of both forward and reverse reads. Low-quality bases were trimmed from the 3’ ends prior to assembly of forward and reverse reads with FLASH^61^. The remaining 66,524,451 high-quality merged reads were clustered with simultaneous chimera removal using UPARSE^62^. After clustering, 37,867,921 read counts mapped to operational taxonomic units (OTUs) at 97% identity (Supplemental Table 6). Taxonomies were assigned to each OTU using the RDP Naïve Bayesian Classifier with custom reference databases^63^. For the 16S rRNA V3-V4 data, this database was compiled from the May 2013 version of the GreenGenes 16S database v13, trimmed to the V3-V4 region. After taxonomies were assigned to each OTU, OTUs were discarded if they were not assigned a Kingdom level RDP classification score of at least 0.5, or if they were not assigned to Kingdom Bacteria, which yielded 10,006 OTUs. In the downstream analyses, we removed low abundance OTUs because in many cases they are artifacts generated through the sequencing process. Samples with low read counts were also removed. To account for differences in sequencing read depth across samples, all samples were normalized to an even read depth of reads per sample random subsampling for specific analyses, or alternatively, by dividing the reads per OTU in a sample by the sum of usable reads in that sample, resulting in a table of relative abundance frequencies.

### Estimates of broad sense heritability of OTU abundance in rhizosphere

To calculate the broad-sense heritability (H^2^) for individual OTU abundances, we fitted the following linear mixed model to OTU abundances of each individual OTU (n=1,189) following a cumulative sum scaling^64^ normalization procedure that adjusted for differences in sequencing depth and fit a normal distribution:

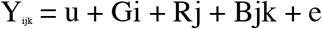

In this model for a given OTU, Y_ijk_ denotes the OTU abundance of the i^th^ genotype evaluated in the kth block of the jth replicate; u denotes the overall mean; Gi is the random effect of the ith genotype; Rj is the random effect of the jth replicate; Bjk is the random effect of the kth block nested within the jth replicate; e denotes the residual error. To account for the spatial effects in the field, additional spatial variables were fitted as random effects using 2-dimensional splines in the above model using an R add-on package “sommer”^65^. H^2^ was estimated as the amount of variance explained by the genotype term (V_G_) relative to the total variance (V_G_ + V_E_/j). Here j is the number of replications. To get the null distribution of H^2^, each OTU was randomly shuffled 1,000 times and then fitted to the same model as described above. Permutation p-value was calculated as the probability of the permuted H^2^ values bigger than the observed H^2^ value.

### Comparative analysis of heritable taxa between sorghum and maize datasets

To identify the degree to which heritable taxa were shared between maize and sorghum, we compared the top 100 most heritable OTUs from both maize datasets (referred to as NAM 2010 and NAM 2015) and the sorghum dataset generated in this study, resulting in a combined dataset of 300 OTUs spanning 65 bacterial orders. As these three experiments were conducted at different field sites, a subset of the orders (n=18) containing heritable OTUs in the maize dataset were not detected in either the heritable or non-heritable fractions of the sorghum dataset and were excluded from subsequent comparative analyses. Of the remaining bacterial orders represented by these heritable OTUs, we determined the number (n=26) that contained heritable OTUs in at least two of the datasets, and the number (n=15) that contained heritable OTUs in all three datasets (Figure 3a). To evaluate whether the degree of overlap in heritable lineages is greater than what would be expected by chance, we performed a permutation test (n=10,000) in which we resampled 100 random OTUs from the 1,189 total sorghum OTUs and recomputed intersections with the two maize datasets. P-values are reported as the number of instances that these permutations returned a greater degree of overlap in these permutations divided by total number of permutations.

### GWAS

For each OTU, GWAS was conducted separately using the best linear unbiased predictors (BLUPs) obtained from the linear mixed model. Population structure was accounted for using statistical methods that allow us to detect both population structure (Q) and relative kinship (K) to control spurious association. The Q model (y = Sα + Qν + e), the K model (y = Sα + Zu + e), and the Q + K model (y = Xβ + Sα + Qν + Zu + e) described previously^66^, were used in our study. In the model equations, y is a vector of phenotypic observation; α is a vector of allelic effects; e is a vector of residual effects; ν is a vector of population effects; β is a vector of fixed effects other than allelic or population group effects; u is a vector of polygenic background effects; Q is the matrix relating y to ν; and X, S, and Z are incidence matrices of 1s and 0s relating y to β, α, and u, respectively. To account for the population structure and genetic relatedness, the first three principal components (PCs) and kinship matrix were calculated using the SNPs obtained from^21^ and fitted into the MLM-based GWAS pipeline for each OTU using GEMMA^67^.

### GWAS validation experiment

For the GWAS validation experiment, the 378 genotypes of the SAP were first subset into lines containing the major (n=343) and minor (n=14) allele for the two haplotypes found at the peak on chromosome 4 described in the text. Including the 178 genotypes not selected for the GWAS, a total of nine sorghum genotypes belonging to the minor allele were selected, with an effort to include genotypes spanning the phylogenetic tree. For each of these nine minor allele lines, another genotype containing the major allele with close overall genetic relatedness was selected, resulting in nine major and nine minor allele containing lines. Two replicates of each line were grown in growth chambers (33°C/28°C, 16h light/ 8h dark, 60% humidity) in a 10% vermiculite/ 90% calcined clay mixture rinsed with a soil wash prepared from a 2:1 ratio of field soil to water from the field site used in the GWAS. Plants were watered daily with approximately 5 ml of autoclaved Milli-Q water using a spray bottle for the first three days, followed by top watering with 15 ml of water every three days. An additional misting was performed to the soil surface every 24 hrs to prevent drying. Following two weeks of growth, plants were harvested and rhizosphere microbiomes extracted as described for the field experiment.

### Microbiome statistical analyses

All statistical analyses of the amplicon datasets were performed in R using the normalized reduced dataset, unless stated otherwise. For alpha-diversity measurement, Shannon’s Diversity was calculated as e^X^, where X is Shannon’s Entropy as determined with the diversity function in the R package vegan^68^. Principal coordinate analyses were performed with the function pcoa in the R package ape^69^, using the Bray-Curtis distance obtained from function vegdist in the R package vegan^68^. Mantel’s tests were used to determine the correlation between host phylogenetic distances and microbiome distances using the mantel function in the R package vegan^68^ with 9,999 permutations, and using Spearman’s correlations to reduce the effect of outliers. Indicator species analyses were performed using the function indval in the R package labdsv^70^, with p-values based on permutation tests run with 10,000 permutations. To account for multiple testing performed for all 430 genera in our dataset, multiple testing correction was performed with an FDR of 0.05 using the p.adjust function in the base R package stats. Canonical Analysis of Principal Coordinates (CAP) was performed for the final validation experiment to test the amount of variance explained by genotypic group using the capscale function in the R package vegan^68^; an ANOVA like permutation test using the sum of all constrained eigenvalues was performed to determine the percent variance explained by each factor using the function anova.cca in the R package vegan^68^.

### Analysis of sorghum RNA-seq datasets

Publicly available sorghum RNA-Seq data for 27 annotated genes in the 1.15 Mb interval of chromosome 4 (Sobic.004G153000 - Sobic.004G155900), were downloaded from phytozome v12.1^25^ (Figure 4c, Supplemental Table 5). Expression datasets were broadly grouped based on the tissue-type from which they were derived (root, leaf, or reproductive). To aid in the visualization of tissue specific expression of genes exhibiting large differences in absolute levels of gene expression, we normalized the Fragments Per Kilobase of transcript per Million mapped reads (FPKM) values for each gene in each tissue type by dividing by the average value of gene expression for that gene across all tissue types. We defined root-specific expression as genes that had a normalized FPKM less than 1 in no more than two root datasets, and a normalized FPKM greater than 1 in no more than two datasets of other tissue types (Figure 4c, Supplemental Table 5).

## Data availability

All datasets and scripts for analysis are available through github (https://github.com/colemanderr-lab/Deng-2020) and all short read data has been submitted to the NCBI SRA.

## Acknowledgments

We thank Dr. Sam Leiboff, Dr. Ling Xu, Edi Wipf, and Tuesday Simmons for their helpful discussions and critical readings of the manuscript. This research was funded by a grant from the US Department of Agriculture (2030-12210-002-00D).

## Author contributions

S.D. conceived and designed the experiments, performed the experiments, analyzed the data, and prepared figures and/or tables; D.C. conceived and designed the experiments, analyzed the data, and prepared figures and/or tables; J.Y. conceived and designed the experiments, and analyzed the data; L.D. performed the experiments; L.W. performed the experiments and analyzed the data; D.C-D. conceived and designed the experiments, analyzed the data, and prepared figures and/or tables; All authors authored or reviewed drafts of the paper and approved the final draft.

