## Supplementary figures and images for "Genome wide association study reveals plant loci controlling heritability of the rhizosphere microbiome"

### Supplemental Figure 1

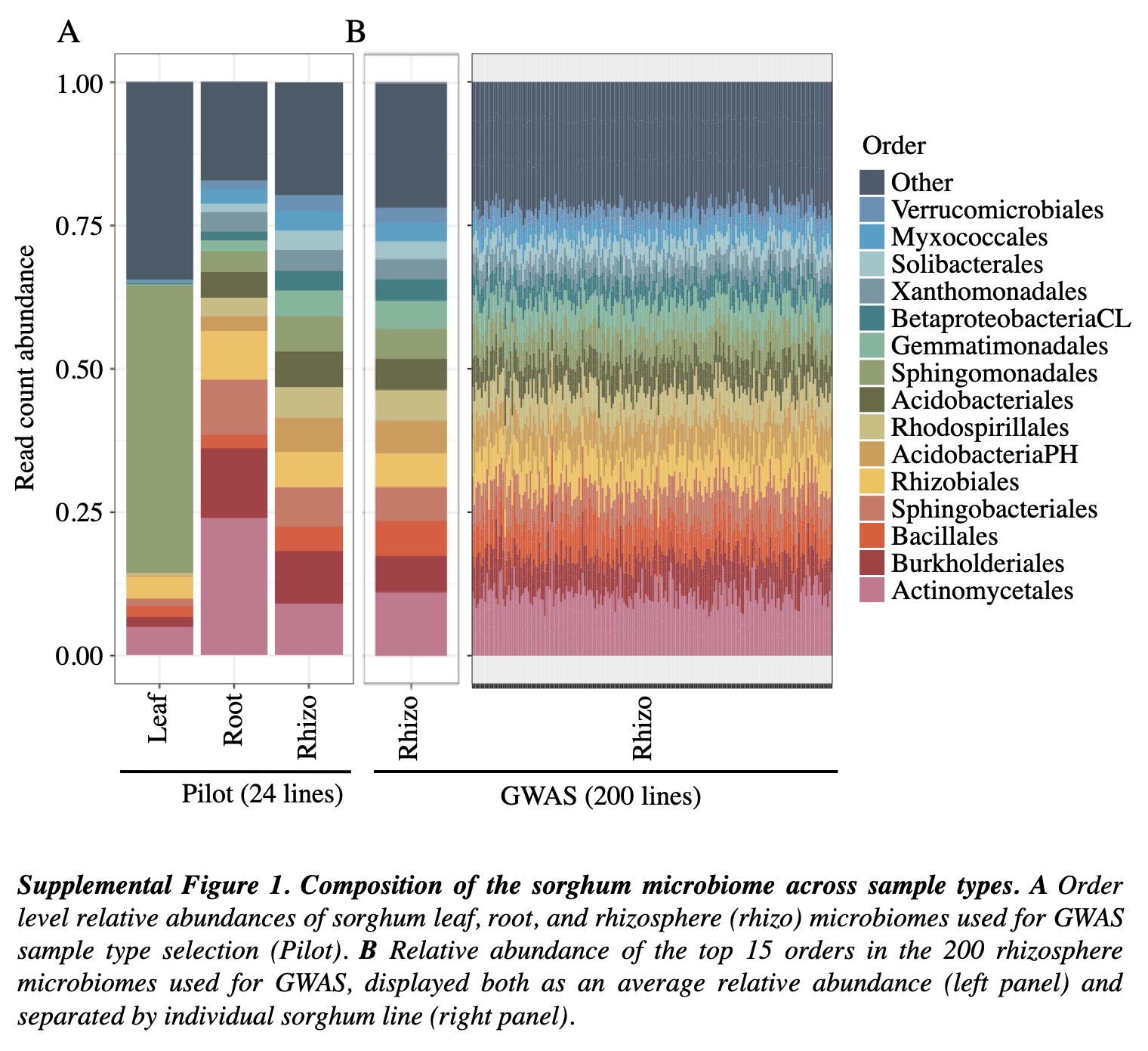

### Supplemental Figure 2

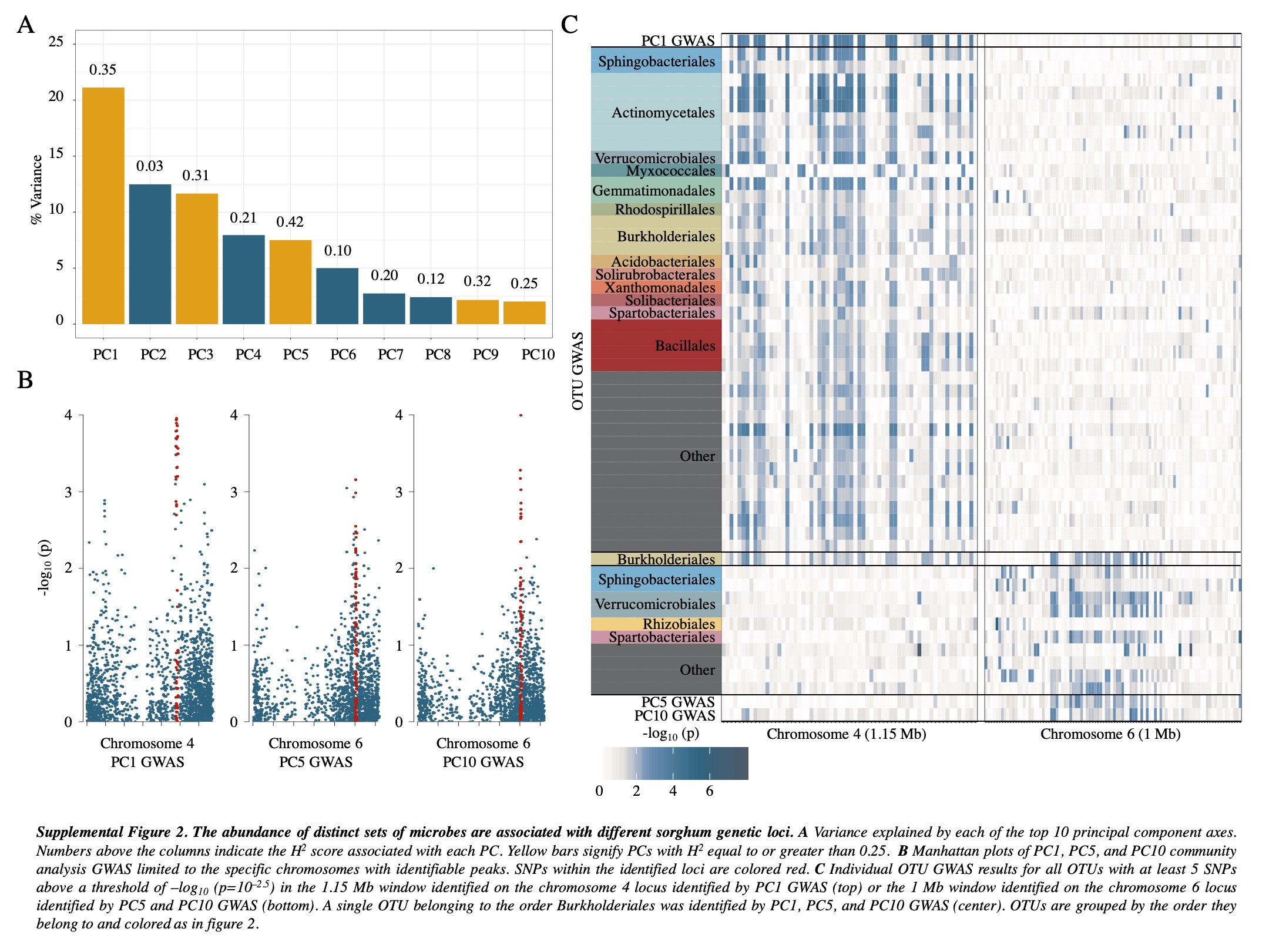

### Supplemental Figure 3

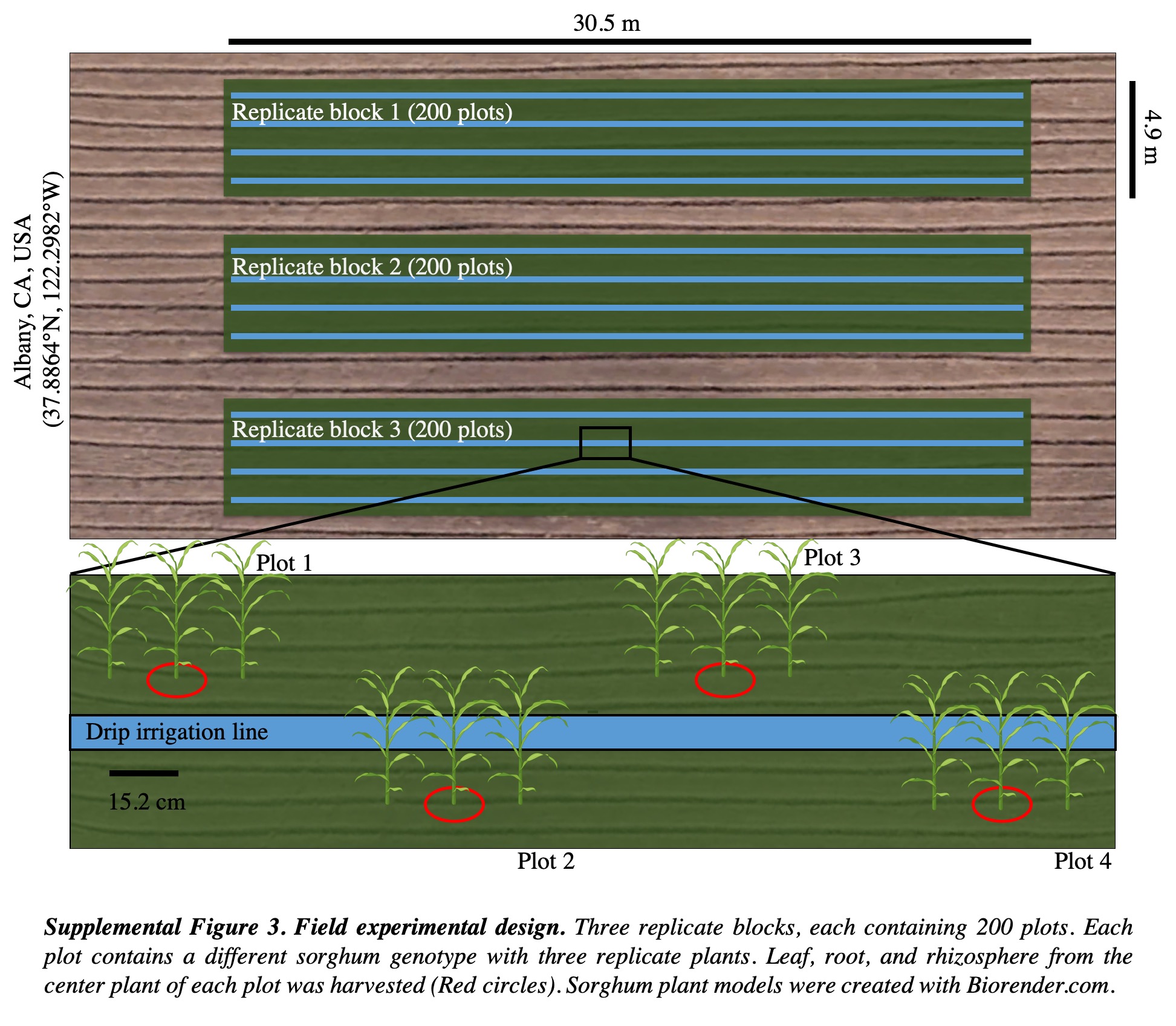
